# Primary productivity declines when species composition and climate are mismatched

**DOI:** 10.64898/2026.05.20.726661

**Authors:** Michael Stemkovski, Kyra Clark-Wolf, Laura E. Dee, Kara C. Dobson, Andrew J. Felton, Thiago Gonçalves-Souza, Giles Hooker, Mevin B. Hooten, Loretta C. Johnson, Mark Morales, Brooke B. Osborne, Malin L. Pinsky, Peter B. Reich, Christine R. Rollinson, Yiluan Song, Nicole K. Ward, Kai Zhu, Peter B. Adler

## Abstract

Climate change drives shifts in species composition, but turnover in many communities lags behind the current pace of change. Anticipating the impact of the resulting community-climate disequilibria on ecosystem functioning is critical. Present-day communities may already be out of equilibrium with climate, providing an opportunity to estimate the effects of disequilibrium before they become more widespread. We analyzed plant community composition and function data from ∼60,000 rangeland monitoring sites across the western US to measure how community-climate disequilibrium contributes to spatial and temporal variation in net primary productivity (NPP) – a key ecosystem function. We found that communities were already substantially out of equilibrium with climate and accounting for this disequilibrium helped explain patterns of NPP. Communities farthest from equilibrium were less productive than those that were closely matched with climate. Our findings suggest that future increases in community-climate disequilibrium may further impair ecosystem functioning.

## Introduction

Climate change is altering species composition in many communities through changes in abundance, immigration, and extirpation. The rate of community change can be quantified using community climate indices, such as the community temperature and precipitation indices (CTI and CPI), which are abundance-weighted averages of species’ realized climate niche optima (Blonder *et al*. 2015; Devictor *et al*. 2008, 2012). Analyses of CTI and CPI time series have shown that communities of short-lived and mobile species can change rapidly, especially when climate shifts are large (Bertrand *et al*. 2011; Pinsky *et al*. 2013; Zhu *et al*. 2024), but many communities are changing slowly, lagging behind climate change (Aguirre-Gutiérrez *et al*. 2025; Alexander *et al*. 2018; Rosenblad *et al*. 2023; Zhu *et al*. 2012). Organisms in these communities will more frequently experience conditions far from their climate niche optima. Mismatch between species composition and the climate – *community-climate disequilibrium –* is likely to grow in ecosystems where climate change outpaces community turnover (Stemkovski *et al*. 2025; Svenning & Sandel 2013). However, the potential consequences of community-climate disequilibrium for ecosystem functioning are unknown.

Large community-climate disequilibria could reduce ecosystem functions such as net primary productivity (NPP) by subjecting plants to climate conditions unfavorable for performance (Stemkovski *et al*. 2026; Svenning & Sandel 2013), suppressing growth, survival, and reproduction, and limiting plants’ ability to convert resources into biomass (Aubree *et al*. 2020; Carroll *et al*. 2023; Lynn *et al*. 2023; Pagel *et al*. 2020). We do not have to wait for climate change to create large community-climate disequilibria in order to test this hypothesis. Contemporary communities exist in various degrees of disequilibrium due to historical climate legacies, non-climatic constraints on species ranges, and tradeoffs between climate tolerance and competitive ability (Gaüzère *et al*. 2018; Liang *et al*. 2018; Pellissier *et al*. 2018; Svenning *et al*. 2015). Comparisons of community climate indices and regional climate have shown that disequilibria are often geographically structured (Bertrand *et al*. 2011; Blonder *et al*. 2015). This existing variation in disequilibria provides an opportunity to measure the impact of disequilibria on NPP before climate change induces larger and more widespread mismatches.

Anticipating how disequilibria will impact NPP, the foundation of food webs and a critical component of the carbon cycle, is critical for informing conservation, management, and nature-based carbon sequestration policies (Bradford *et al*. 2018; Chapin *et al*. 1997; Friedlingstein *et al*. 2022; Schuurman *et al*. 2022). In water-limited ecosystems, mean annual precipitation (MAP) is a strong predictor of NPP: productivity is highest in the wettest sites and years (Huxman *et al*. 2004; Knapp & Smith 2001; Sala *et al*. 1988). Mean annual temperature (MAT) also influences NPP, though the direction of its effect varies across dryland vegetation types (Felton *et al*. 2022). These results are based on statistical models that predict NPP as a function of climate variables, with little or no consideration of species composition (e.g., Felton *et al*. 2021). If community-climate disequilibria do influence NPP, then including information about them should improve these models, explaining more variation in historical patterns of NPP across space and time and potentially providing more skillful forecasts of climate-driven changes in NPP.

We tested three hypotheses about community-climate disequilibrium and NPP in drylands of the western United States. 1) Dryland plant communities are not in equilibrium with climate, particularly with respect to water availability, because these communities are composed of species at the dry edge of their ranges. To test this hypothesis, we described community climate niches using relative abundance data from ∼60,000 rangeland plant surveys (US BLM 2025) and North American occurrence records (GBIF.org 2025). 2) Community climate niches can help explain variation in NPP because communities function differently depending on the magnitude of their disequilibria. We tested this hypothesis by comparing a suite of models that explain spatiotemporal variation in remotely-sensed NPP using either climate anomalies centered on long-term climatological averages or climate disequilibria centered on community climate niches. 3) Positive and negative disequilibria reduce NPP because they subject species to conditions outside of their climate tolerance (Figure 1). To test this hypothesis, we draw inference from a model relating variation in NPP to the magnitude of precipitation and temperature disequilibria across space and time.

**Figure 1.**
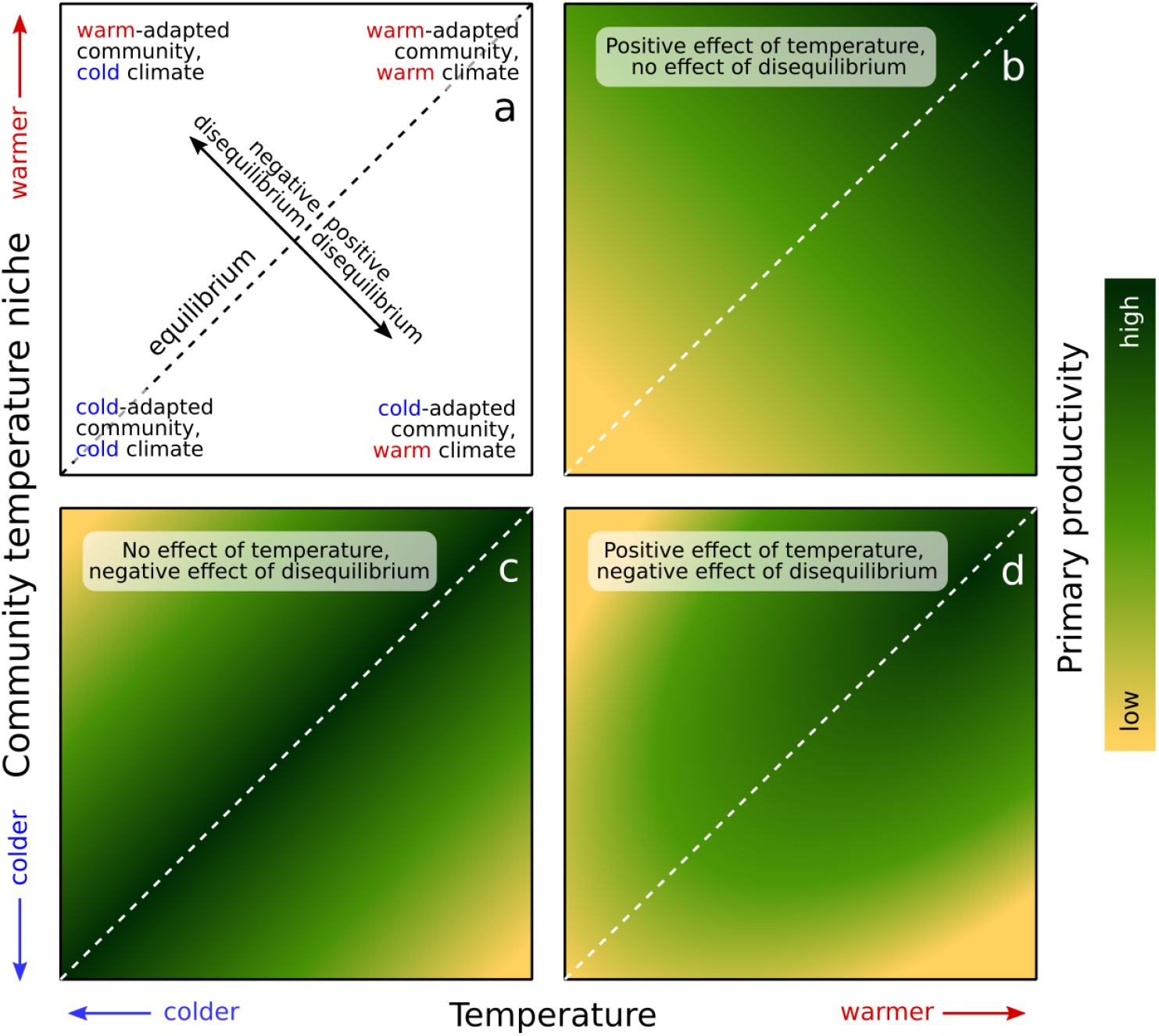
Primary productivity depends directly on climate and may be affected by community-climate disequilibrium. Panel a: In the example of temperature, communities might be suited to colder climate than they experience (negative disequilibrium; top left), warmer climate (positive disequilibrium; bottom right), or the same climate (equilibrium; dashed diagonal line). Communities depart from equilibrium due to changes in climate (left-right), mismatched community composition (up-down), or a combination of both (perpendicular to equilibrium line). Panels b-d represent different hypotheses about the interplay of direct climate sensitivity and the effects of community-climate disequilibrium on NPP.

### Rangeland plant communities are not in equilibrium with climate

Disequilibrium between species composition and the climate was widespread across western US drylands. 72% of sites in the study area had climate colder than the community climate niche, and 86% had climates too dry for the community. On average, CPI and CTI were 93 mm and 0.95 °C lower than MAP and MAT, respectively. This bias indicates that most rangeland communities in the study area disproportionately comprised species at the cold and dry edge of their ranges. Disequilibria – which are calculated in relation to long-term climatological averages (MAP-CPI and MAT-CTI) – were of a similar magnitude to interannual climate variation experienced by communities. On average across all sites in the study area, 57% of sites had community climate niches more than 1°C colder or hotter than the site’s mean annual temperature, and 30% were more than 2°C out of equilibrium (Figure 2a). 49% of sites were composed of species whose averaged climate niche optima were more than 100mm wetter or drier than the mean annual precipitation in that area, and 10% were more than 200mm wetter or drier (Figure 2b). By comparison, annual temperatures at sites varied by 2.4°C on average (2SD of interannual temperature distribution), and annual precipitation varied by 332.9mm. This widespread, biased disequilibrium suggests that future climate warming (in the absence of community turnover) would bring communities closer to most species’ realized niche optima, but areas in which precipitation will decrease would be more heavily impacted than if those communities were in equilibrium before the onset of climate change. Though the precise causes of disequilibria cannot be ascertained with the present data, other research points to historical climate legacies (Svenning et al. 2015), lags in dispersal, establishment, and extirpation (Alexander et al. 2018), and non-climatic drivers of community composition such as disturbance (Johnstone et al. 2016) and biotic interactions (Wisz et al. 2013).

**Figure 2.**
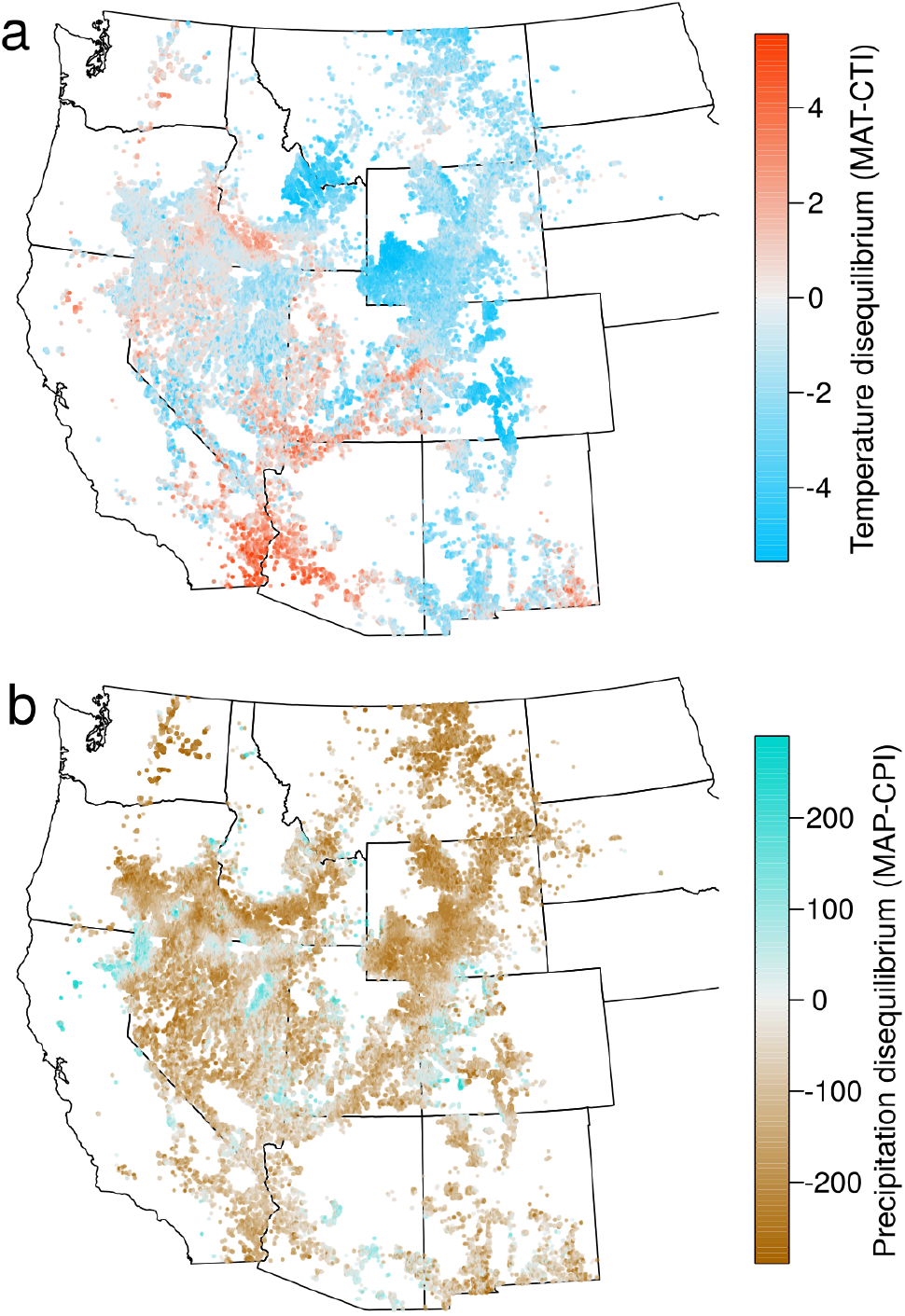
Community-climate disequilibria were widespread and geographically structured. Panel a: most sites across the study region showed negative temperature disequilibria, with climate “too cold” (blue) for the community, but many also experienced conditions that were “too warm” (red). Panel b: most sites showed negative precipitation disequilibria, with climate “too dry” (brown) for the community.

Disequilibria varied substantially across space. Precipitation disequilibrium was negative across most ecoregions, but was especially pronounced in the Northwest Great/Glaciated Plains and the Snake River Plain (Supplement 2). Temperature disequilibrium was most negative in the Middle and Southern Rockies, but was positive in the Sonoran Basin and Range. The Mojave and Northern Basin and Range ecoregions were nearest to equilibrium in terms of temperature. Regional climate explained a moderate amount of this spatial variation in disequilibria: Warmer and wetter regions tended to have positive community temperature and precipitation disequilibria, with moderately strong correlations between MAP and precipitation disequilibrium (r=0.49) and MAT and temperature disequilibrium (r=0.59). The greatest absolute disequilibria were observed in areas with the most extreme climates (Figure S1.2). This pattern may reflect source-sink dynamics in community assembly, in which dispersal of propagules from neighboring communities counteracts filtering of species by the climate (Gravel *et al*. 2010).

While the lower dispersion of community climate niches compared to sites’ climate may be, in part, a byproduct of community climate index calculation because the method averages across species (Lepš *et al*. 2011), it is unlikely to be a result of artificial niche truncation because species climate niches were calculated using independent occurrence records across North America (Anselmetto *et al*. 2025).

### Community climate niches help explain variation in net primary productivity

In spatiotemporal analyses of NPP, interannual climate variation is often characterized as anomalously hot/cold and wet/dry with respect to a long-term climatological average, typically from the second half of the 20th century (e.g., Felton *et al*. 2021; Mohamed *et al*. 2004; Niu *et al*. 2017). Using annual anomalies in this way effectively assumes that communities were in equilibrium before the onset of recent climate change (Evans *et al*. 2025). To test whether community-climate niches are more ecologically relevant baselines than climatological averages (Figure S1.3), we compared the predictive ability of a suite of models built around mean annual precipitation (MAP) and temperature (MAT) against a complementary set using community precipitation (CPI) and temperature indices (CTI).

Accounting for community-climate disequilibrium by using community climate niches in place of climatological averages helped explain variation in NPP (Figure S1.4a). The best-supported model used community climate niches as a baseline in all fixed effect terms and allowed for nonlinear effects of disequilibria:

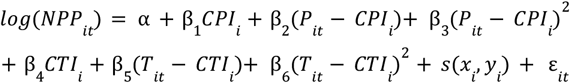

In this model, site (*i*) average NPP is predicted by community climate indices (*CPI* & *CTI*), and interannual (*t*) variation in NPP is predicted by annual disequilibria relative to the community climate indices (*P-CPI* & *T-CTI*). This model explained 51% of the variation in NPP before accounting for spatial autocorrelation, and 63% with a spatial smooth random effect *s*. In contrast, treating communities as if they are in equilibrium by calculating annual climate anomalies against MAP and MAT (Figure S1.4, eq. 3) explained 44% of variation in NPP, and 62% after accounting for spatial autocorrelation. The CPI & CTI model also made more accurate out-of-sample predictions when data were split into independent training and test sets (Figure S1.4b). These results run counter to the widespread, often implicit assumption that climate in the twentieth century is an accurate baseline against which to judge ecological responses to recent climate change. Our approach demonstrates that such assumptions are not necessary when community composition data are available.

The results of the model comparison are likely conservative for at least two reasons. First, measuring community climate niches using species occurrence data is a crude approach.

Not least among its problems is the assumption that species geographic ranges track their climate niches in space without lags (Essl *et al*. 2023; Lalechère *et al*. 2025). Second, we do not attempt to account for local adaptation. Direct measurements of species climate performance curves in-situ (Rezende & Bozinovic 2019) or locally-tuned species climate index estimates (Zhu *et al*. 2024) could further clarify how close individual populations are to their optima. Both of these problems likely contribute noise to our measures of community climate indices, which would make it less probable that the indices would explain variation in NPP, unless the signal of disequilibrium is truly strong. It may also be that community climate indices provide information about on-the-ground microclimate conditions that are missed by gridded macroclimate data; However, this should not affect our conclusion about their predictive power because the climate and NPP data have similar spatial resolution.

### Net primary productivity was reduced under large disequilibria

Communities with wetter and colder climate niches were more productive than drier and warmer communities (Figure 4a,c). These patterns reflect the direct effect of climate on NPP, which might be expected when communities are in equilibrium with climate (dashed white lines in Figure 4e,f). In other words, if community composition were allowed to acclimate to climate for a long period of time, this would be the expected relationship between NPP and MAP and MAT in the study region. This finding agrees with macro-scale observations of higher productivity under lower water stress (Huxman *et al*. 2004; Maurer *et al*. 2020). This direct relationship between climate and productivity must be accounted for in order to understand the effect of disequilibrium.

After accounting for the effects of CPI and CTI, we found that productivity was lower when disequilibria were large. Overall, NPP was greatest when precipitation and temperature experienced by communities were close to matching to their CPI and CTI values (Figure 4b,d). Both large negative (climate drier or colder than community niche) and positive (climate wetter and warmer than community niche) disequilibria were associated with decreased NPP. Large community-climate disequilibria likely subject species to conditions that suppress growth, survival, and reproduction, limiting their ability to convert available resources into biomass (Lynn et al. 2023; Pagel et al. 2020). However, the relationship was asymmetrical: NPP was maximized under moderate negative precipitation disequilibrium (-271 mm) and temperature disequilibria (-3.6 °C). This asymmetry may be related to the earlier finding of negative precipitation and temperature disequilibria across the study region (Figure 2). CTI and CPI could be biased measures of community climate niches in this region, or the same ecological processes that maintain disequilibria might also maximize productivity.

The effects of disequilibrium were negative in nearly every ecoregion (Supplement 2), though the levels at which NPP was maximized varied substantially, and the direct relationships between CTI and NPP were idiosyncratic (Figure S2.3-2.4). However, parameter estimates were not greatly influenced by any single ecoregion (Figure S2.5-2.6), indicating that our overall results are robust across ecological contexts. Many landscape and land-use variables covary with climate and NPP, but they would confound our inference only if they also covaried with community-climate disequilibrium – a more specific relationship for which we lack strong *a priori* expectations. The finding of a negative effect of disequilibrium was corroborated by a complementary modeling approach that isolates the influence of disequilibrium into a single parameter and allows for a nonlinear relationship between MAP and NPP (Supplement 3).

### Implications

Variation in community-climate disequilibrium across communities provides insights about the functioning of communities in the future. We found that extreme disequilibria were associated with reduced productivity, suggesting that future disequilibria will also hamper ecosystem functions. Our findings suggest that increases in temperature will decrease productivity in most communities, with the exception of some warm-adapted communities where warming might reduce disequilibrium and ameliorate the negative direct effects of warming. Climate change is forecasted to have geographically heterogeneous effects on precipitation across the study area (IPCC 2023). Our results suggest that many communities are pre-adapted for increasing moisture, but the resulting boost in productivity might be less than expected due to the effect of disequilibrium; Loss of precipitation is unlikely to increase productivity in any communities. While the magnitude of present-day disequilibrium is comparable to interannual climate variation and extreme disequilibria were rare in the study system, ongoing climate change has the potential to produce more widespread disequilibria if it outpaces species dispersal, establishment, and growth. The rate at which community composition will turn over in response to climate change is, therefore, a critical uncertainty in forecasts of primary productivity (Felton *et al*. 2022). We found evidence that warm- and wet-adapted species have increased in relative abundance across survey years, but repeated surveys of communities are needed to robustly quantify the pace of community climate niche shifts relative to the pace of climate change (Supplement 4.1). Future research should focus on the timescales of community acclimation and on the sensitivity of various ecosystem functions to disequilibrium (Stemkovski *et al*. 2026).

Our findings highlight a potential trade-off between land management to preserve historical species assemblages and management to maintain ecosystem function. Efforts to resist compositional transitions may intensify community-climate disequilibrium in the future, potentially reducing primary productivity and the ecosystem services that productivity supports (Bradford *et al*. 2018; Schuurman *et al*. 2022). Conversely, management to accelerate ecological acclimation through assisted migration may be beneficial when maximizing productivity is a primary objective (Hällfors *et al*. 2017). Our measurements of present-day disequilibria may be helpful for informing such management decisions (Stemkovski *et al*. 2025).

We have shown a novel application of community climate indices to predict ecosystem function. Ecologists have long sought to make sense of the complexity of communities by summarizing them with synthetic metrics such as species diversity, richness, and evenness, measures of phylogenetic and interaction network structure, and community-weighted functional traits (Cadotte 2017; Cavender‐Bares *et al*. 2009; Harvey *et al*. 2017). Much research effort has gone into investigating the relationship between biodiversity and ecosystem function (Tilman *et al*. 2014). Community climate indices offer a way to investigate the biodiversity-ecosystem function relationship that is particularly relevant in the context of climate change; these indices summarize communities in a way that makes them directly comparable to climate by expressing the biotic and abiotic environment in the same units. To date, community climate indices have been used almost exclusively to summarize the effects of climate change on community composition, and their potential for explaining ecosystem functioning has not been appreciated. We hope that this study provides a blueprint for new research that draws insights by relaxing equilibrium assumptions. There is huge potential for reanalysis of existing datasets because the requisite components (species relative abundances, climate time series, ecosystem function data) are widely, and increasingly, available (Abatzoglou *et al*. 2018; Dornelas *et al*. 2025; Robinson *et al*. 2018).

## Acknowledgements

This work was funded by NSF BoCP grant 204754. We thank Annie Schiffer, Matthew Peña, and Isabela Pedro for providing helpful advice at earlier stages of this work. John Bradford, Abigail Lynch, and Melissa Pastore contributed significantly to this study.

## Methods

We utilized four data sources to assess community-climate disequilibrium and its effect on productivity in western US rangelands in the following workflow (Figure 4). First, we compiled relative abundance data from on-the-ground surveys from the U.S. Bureau of Land Management’s Assessment and Inventory (AIM) program (US BLM 2025). Second, we extracted North American occurrence records for all species in the surveys from the Global Bioinformatics Facility (GBIF.org 2025). Third, we recorded annual temperature and precipitation data from Daymet at GBIF the occurrence locations and AIM sites. Fifth, we estimated species climate niches as the average climate across their GBIF occurrences. Sixth, we quantified community climate indices and disequilibria at AIM sites by taking abundance-weighted averages of species climate niches. Finally, we extracted remotely sensed NPP data from RAP at the AIM sites and modeled productivity as a function of climate and disequilibrium.

### Community composition data

We quantified plant community-climate disequilibrium at 58,876 sites across the western United States using species composition data from the US Bureau of Land Management’s Assessment Inventory and Monitoring (AIM) program. The AIM dataset spans diverse rangeland biomes, including cold desert shrublands, warm desert grasslands, woodlands, forested mountains, northern Great Plains grasslands, and California mediterranean grasslands. At each ∼40m^2^ site, standard line-point-intercept protocols were used to measure species percent aboveground cover (a proxy for relative abundance) between 2010 and 2023. We assessed only upland sites, excluding riparian areas whose vegetation may depend on precipitation far from the study locations. We accessed AIM data using the *trex* R-package curated by the Landscape Data Commons platform. We used the USDA PLANTS database to translate species codes, further disambiguated species names using the Catalogue of Life, GBIF taxonomic backbone, and Encyclopedia of Life databases through the *taxize* R-package, and excluded any remaining ambiguous percent cover records. We excluded records at the genus level or higher, and we did not distinguish among subspecies. We excluded data from BLM lands in Alaska to avoid major discontinuity across biomes and climate gradients. We were able to use data on 5,083 unique species across all sites, including annual and perennial grasses, forbs, shrubs, and trees.

### Climate data

We used temperature, precipitation, and solar radiation data from Daymet v4 to estimate species climate niche indices, site climate, and community-climate disequilibria (Thornton *et al*. 2022). The Daymet product is based on weather station data and captures for effects of topography – such as rain shadows – in its spatial interpolation procedure. Accounting for complex topography was critical for analyzing the AIM dataset, as many sites were in mountainous terrain far from weather stations. At each AIM site, we extracted annual weather values over the water-year (October 1 of the previous year to September 30 of the given year), and we averaged across all years (1986-2023) to get climatological variables at GBIF occurrence locations (see next section).

### Community climate niches & disequilibrium

We estimated realized climate niche optima for each of the 5,083 species in the dataset by finding the median Daymet value of mean annual temperature (MAT) and precipitation (MAP) across each species’ GBIF occurrence records (GBIF.org 2025). To match the spatial extent of Daymet, we selected occurrences from Canada, the United States, and Mexico, and we thinned the occurrences to 1 per 1km^2^ grid cell to minimize observation bias. We excluded records from islands, fossil specimens, observations dated before 1900, and records with uncertain coordinates.

We then calculated community climate indices (CTI & CPI) at each site in the AIM dataset as the abundance-weighted mean of species realized climate optima (following Devictor *et al*. 2008). This approach expresses communities in units of climate variables, allowing us to calculate the temperature and precipitation disequilibria simply as MAT-CTI (°C) and MAP-CPI (mm). Positive temperature disequilibrium values indicate a community experiencing a climate that is “too hot” for its species, and negative precipitation disequilibrium values indicate a climate that is “too dry” for the community. We calculated annual disequilibrium values as the difference between the community climate niches as and each years’ mean temperature and total precipitation to account for variation in MAT and MAP. We were forced to keep CTI and CPI values constant across years because AIM sites were not resurveyed. The assumption that CTI and CPI stay relatively constant compared to interannual temperature and precipitation variation is crude but not unreasonable because community niche shifts have typically been much slower than climate change, and are not very sensitive to interannual variation. We used variation in AIM survey years (2010-2023) to check the impact of this assumption by modeling CPI and CTI as linear functions of survey year (Supplement 4.1). We focused on temperature and precipitation data because these are the most commonly used climate metrics, and CTI and CPI are the main community climate indices used in other studies.

### NPP data

We obtained annual NPP data from the Rangeland Analysis Platform (RAP) using Google Earth Engine at a 30m spatial resolution at each study site location (Jones *et al*. 2021). RAP estimates total (both above- and below-ground) NPP from downscaled Landsat satellite imagery using a model based on MODIS17. RAP NPP estimates have been validated with independent flux tower measurements and compared against on-the-ground rangeland vegetation survey data. NPP data were available from 1986 to 2023, and we used this time range for the entire analysis.

### Mathematical models

To learn whether accounting for community-climate disequilibrium can help explain temporal and spatial variation in NPP, we assessed a set of linear models with predictor variables for interannually varying weather, long-term climatological averages, and community climate niches (Figure 3). We based our models on classical models used to predict NPP using climate variables, where spatial variation in climate is partitioned across sites, and interannual climate variation is included within sites. In these models, temperature and precipitation weather variables (*T* & *P*) vary across sites *i* and years *t*, and long-term climatology and community climate niches (*MAP, MAT, CPI*, & *CTI*) vary across sites. We started with the simplest models that account only for spatial climate variation, and iteratively increased model complexity to include spatio-temporal covariates and nonlinear terms. We introduced nonlinearity using squared anomaly and disequilibrium terms to reflect the prediction that both extremely high and low climate extremes may affect NPP similarly. For example, NPP might be hampered in a years that is unusually cold and a year that is unusually hot. Climatology and community climate niche variables were compared as alternative baselines against which to compute annual anomalies (*P-MAP* & *T-MAT*) and annual disequilibria (*P-CPI* & *T-CTI*), resulting in two sets of models. This modeling approach captures the effects of disequilibria and tests the hypothesis that community climate indices are a more ecological relevant baseline than climatic averages against which to judge the effects of climate variation. However, the models mix the direct effects of climate on NPP and disequilibrium effects among its parameters; we performed additional modeling to isolate the effect of disequilibrium to its own parameter (Supplement 3).

**Figure 3.**
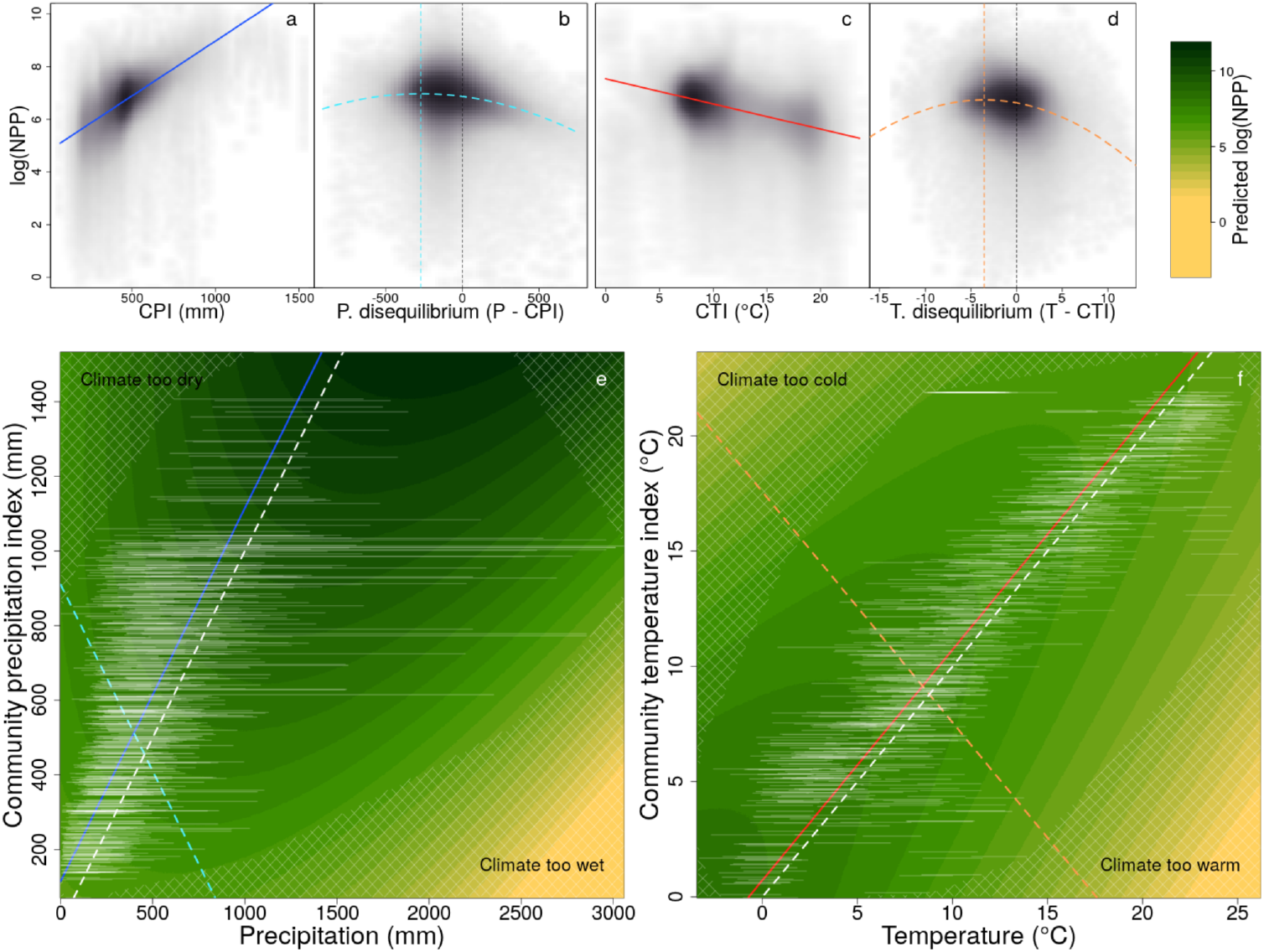
Primary productivity was lowest at sites and years in which community climate niches were mismatched with climate. Panel a-d show one-dimensional conditional predictions from cross-sections of the two-dimensional response surfaces shown in panels e and f. For precipitation (left), panel a shows the effect of community precipitation index (CPI) along a cross-section parallel to the equilibrium line and panel b shows the effect of precipitation disequilibrium along a cross-section perpendicular to the equilibrium line. Panels c-d show the same plots for the temperature variables. In panels a–d, colored curves are model predictions of log(NPP), vertical colored lines show the disequilibria at which log(NPP) was maximized, and gray density clouds show the distribution of partial residuals around the fitted relationships. Panels e and f show predicted log(NPP) across precipitation/CPI space and temperature/CTI space, respectively, with darker green indicating greater predicted productivity. The dashed white one-to-one lines show where community climate indices match climate. Colored solid and dashed lines show the cross-sections plotted in the corresponding top panels. White horizontal lines show the 95% range of interannual climate variation for a sample of 1,000 sites stratified across CPI and CTI variables for visual clarity. Cross-hatched regions indicate predictions extrapolated beyond the support of the observed data.

**Figure 4.**
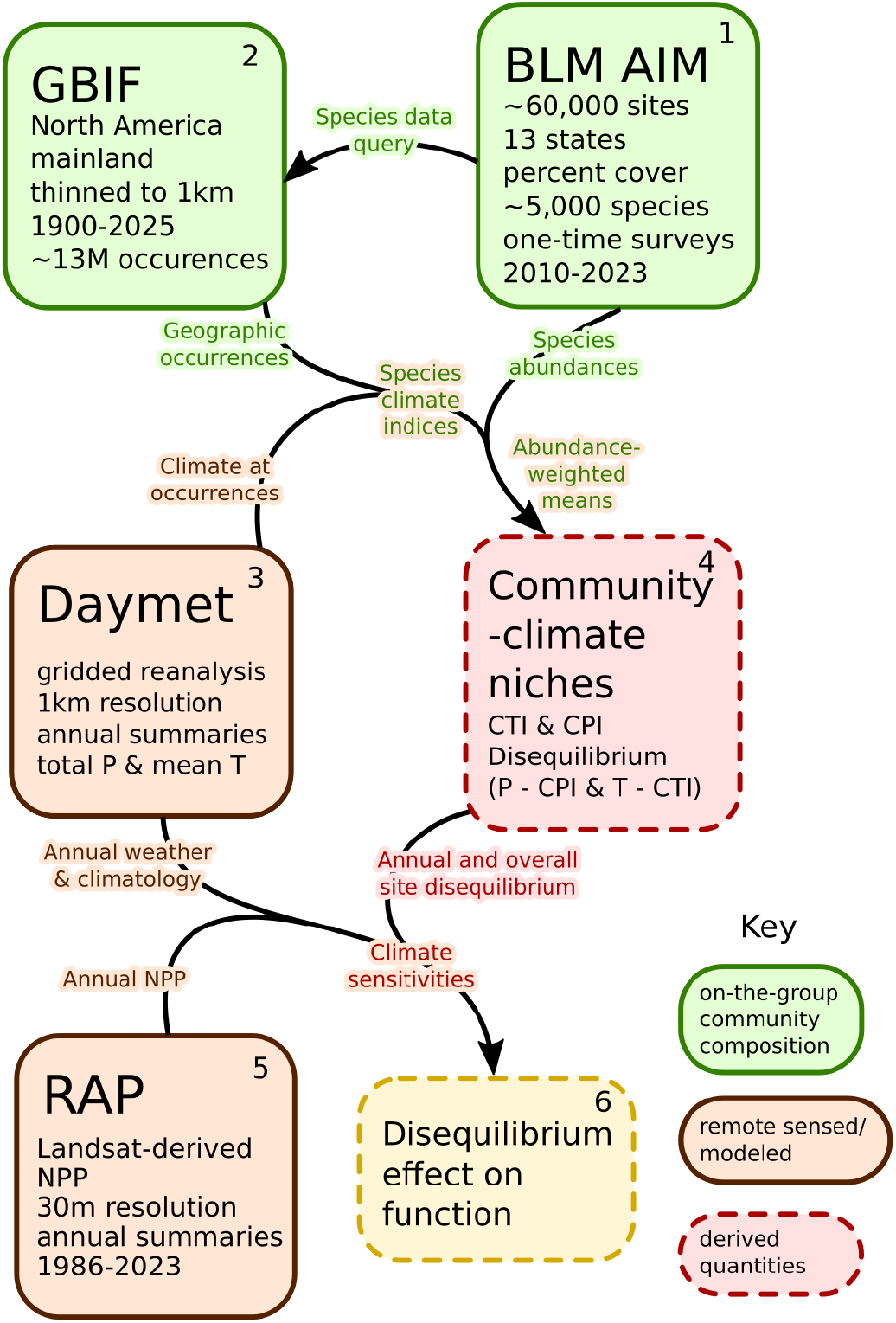
Data analysis workflow.

### Statistical analysis

We compared models using adjusted coefficients of determinations (R^2^) and out-of-sample predictive performance. We evaluated predictive performance using five-fold cross-validation with folds assigned at the site level. Models were fit to training data and evaluated on held-out sites, and performance was summarized as root mean squared error (RMSE) of log(NPP) across folds. We selected the best-supported model (equation 8 from Figure S1.4) to test the effects of community-climate disequilibrium on NPP. To draw inference about the direct effects of climate and effects of disequilibrium on productivity (Stemkovski *et al*. 2026), we examined cross-sections of response surfaces across CPI/precipitation and CTI/temperature variable space. To isolate the direct relationship between climate and productivity, we computed conditional predictions parallel to equilibrium one-to-one lines where community precipitation indices equal precipitation (with an offset to account for overall disequilibrium). To isolate the effects of disequilibria, we computed conditional predictions perpendicular to the equilibrium lines, where a shift in disequilibrium is the product of equal changes in community climate indices and climate variables; this approach avoids conflating the direct effects of climate with the effects of disequilibrium. To understand how accounting for community composition affected inference, we compared the disequilibrium model against the analogous anomaly model (equation 4 from Figure S1.4).

Many AIM study sites are clustered together on patchy BLM lands, and their proximity violates the assumption that they offer independent measures of community composition and productivity. To ensure robust inference, we fit a generalized additive model with a spatial spline over the site coordinates to account for spatial autocorrelation using the *bam* function from the *mgcv* R-package (Wood *et al*. 2015). For the spatial spline, we used a thin-plate smoothing basis with 2nd-order shrinkage to prevent overfitting. We capped basis function dimensionality at 200, used fast-REML to estimate the smoothness parameter, and we discretized the smooth covariates to reduce computation time using default *bam* settings. In R formula syntax, the smooth term was represented this way: *s(x, y, bs = ‘ts’, k=200, m=2)*. The effects of weather and community climate niche variables were modeled without smoothing for interpretable parameter estimates.

We note that drawing inference from either temporal or spatial variation alone (vertically or horizontally in Figure 4e,f) would be inappropriate; holding spatial variation in CPI/CTI constant and looking only at the response across time (or vice-versa) would conflate direct climate effects and disequilibrium effects because both climate and disequilibria vary across years. Geometrically speaking, the effect of disequilibrium runs perpendicular to the equilibrium CPI/P and CTI/T lines (diagonal in Figure 3e,f).

