## Supplementary materials for "Primary productivity declines when species composition and climate are mismatched"

### Supplement 1. Additional figures

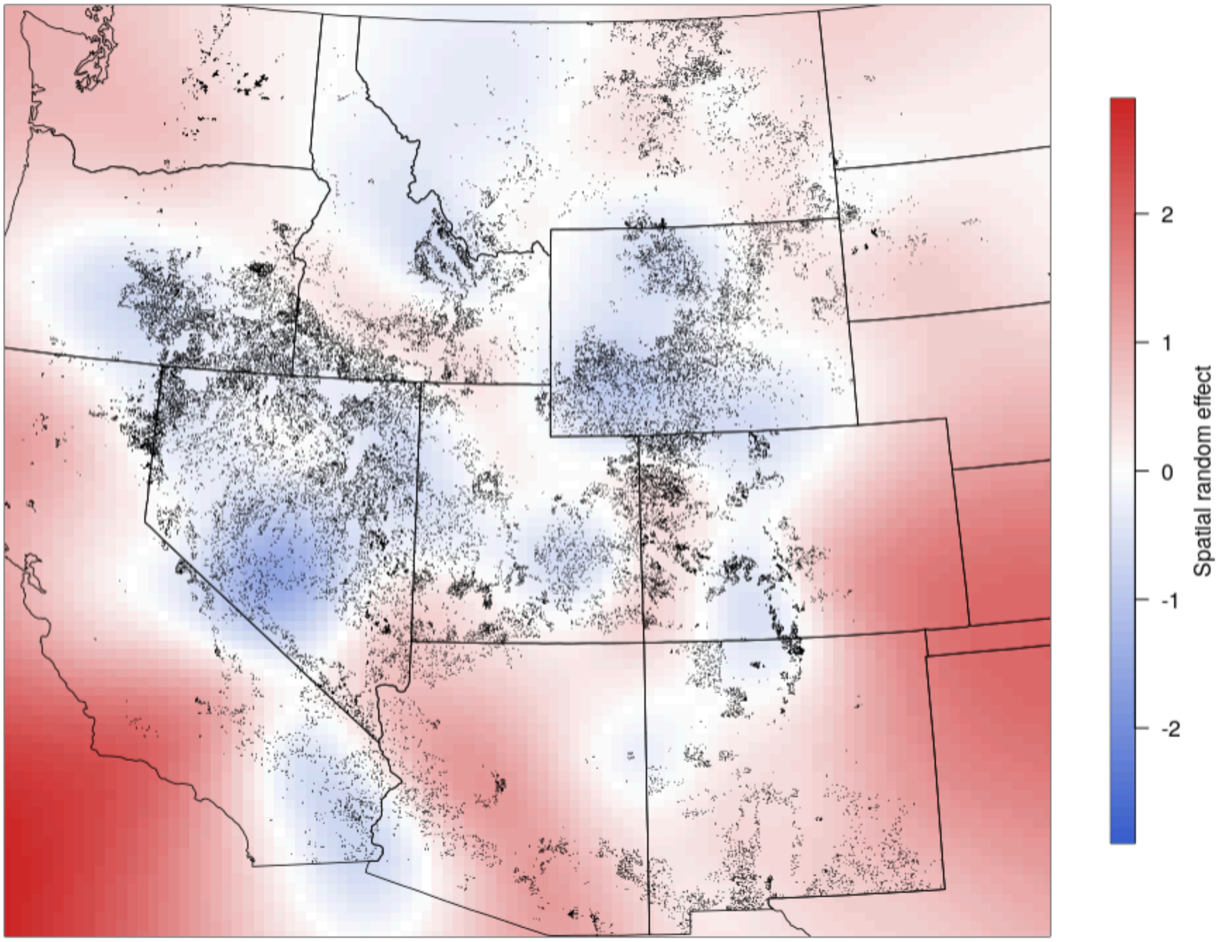


**Figure S1.1.** Spatial random effect from GAM fit, in units of log(NPP).

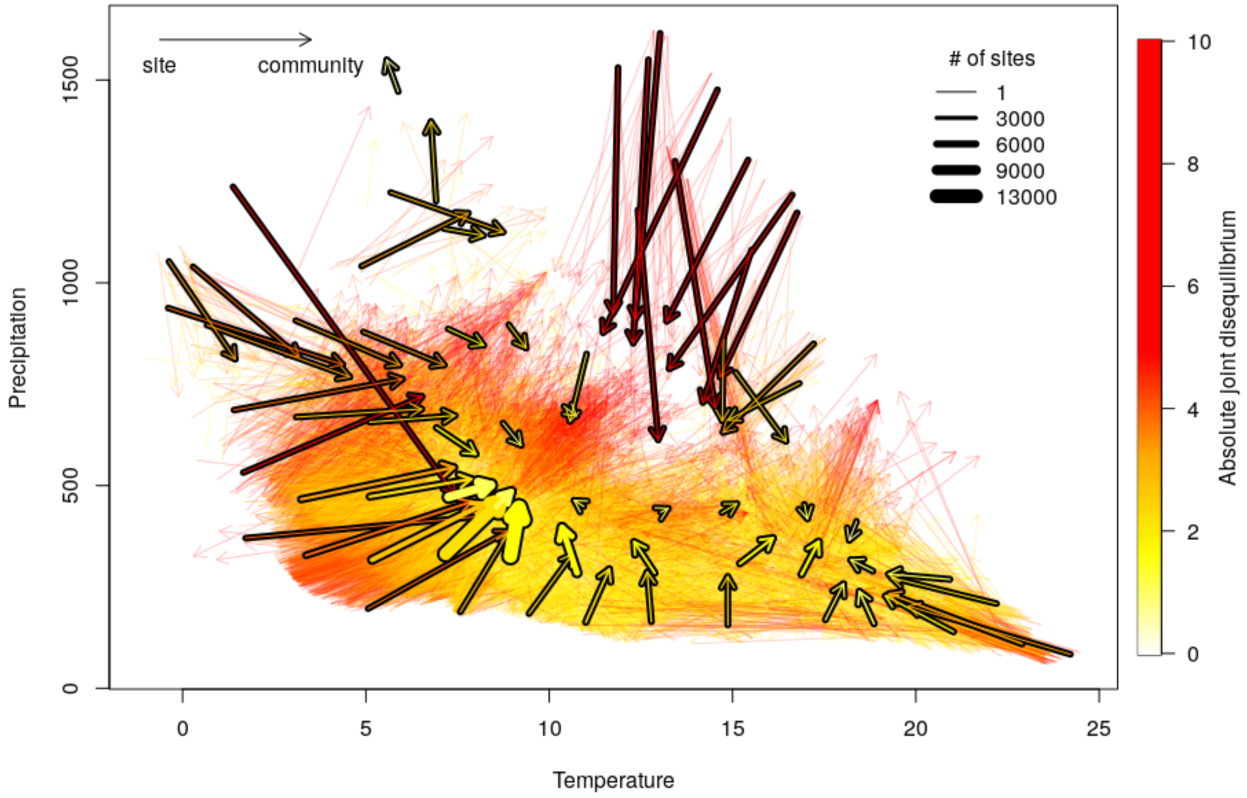


**Figure S1.2**. Summary of community-climate disequilibria across both temperature and precipitation variables. The base of each arrow show sites' MAT and MAP values, and tips of arrows show sites' CTI and CPI values. Arrow colors are show the magnitude of joint disequilibria, with temperature and precipitation disequilibria standardized according to their standard deviations to make the variables comparable. Thin arrows show disequilibria at individual sites. Arrows with black outlines show averages across multiple sites within an aggregating grid across the climate variables. The thickness of the outlined arrows represents the number of sites that make up the average.

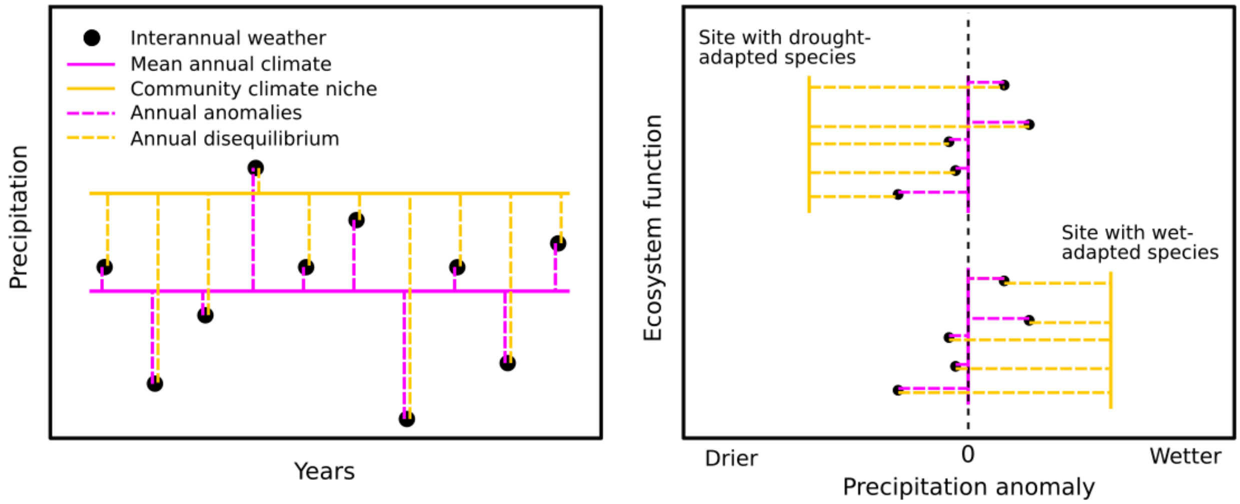


**Figure S1.3.** Left panel: Using long-term climatological average (solid pink line) as a baseline for measuring interannual climate variation necessarily results in a symmetrical spread of positive and negative annual anomalies (pink dashed lines). When annual anomalies are used to predict ecosystem function, it is implicitly assumed that communities experience an equal number of wet and dry years over the long term. Using community climate niches as a baseline against which to partition interannual variation allows for this assumption to be relaxed. In this example, a community climate niche (yellow solid line) that is higher than the mean annual precipitation indicates that most species are suited to wetter climate than what the community experiences. This results in an asymmetric distribution of annual disequilibria, with most years registering as dry years (yellow dashed lines). Right panel: Communities differ in the magnitude and direction of their disequilibria, and this may help explain patterns of ecosystem function. In this example, two sites receive the same precipitation, but differ in their ecosystem function. The site with a drought-adapted species assemblage (top) experiences unexpectedly wet conditions (yellow lines, left) and consequently higher ecosystem function, whereas the site with species adapted for high water availability (bottom) experiences drought conditions (yellow lines, right) and reduced function. Using climatology as a reference (pink lines) would treat the sites as comparable and would fail to explain differences in ecosystem function across the sites.

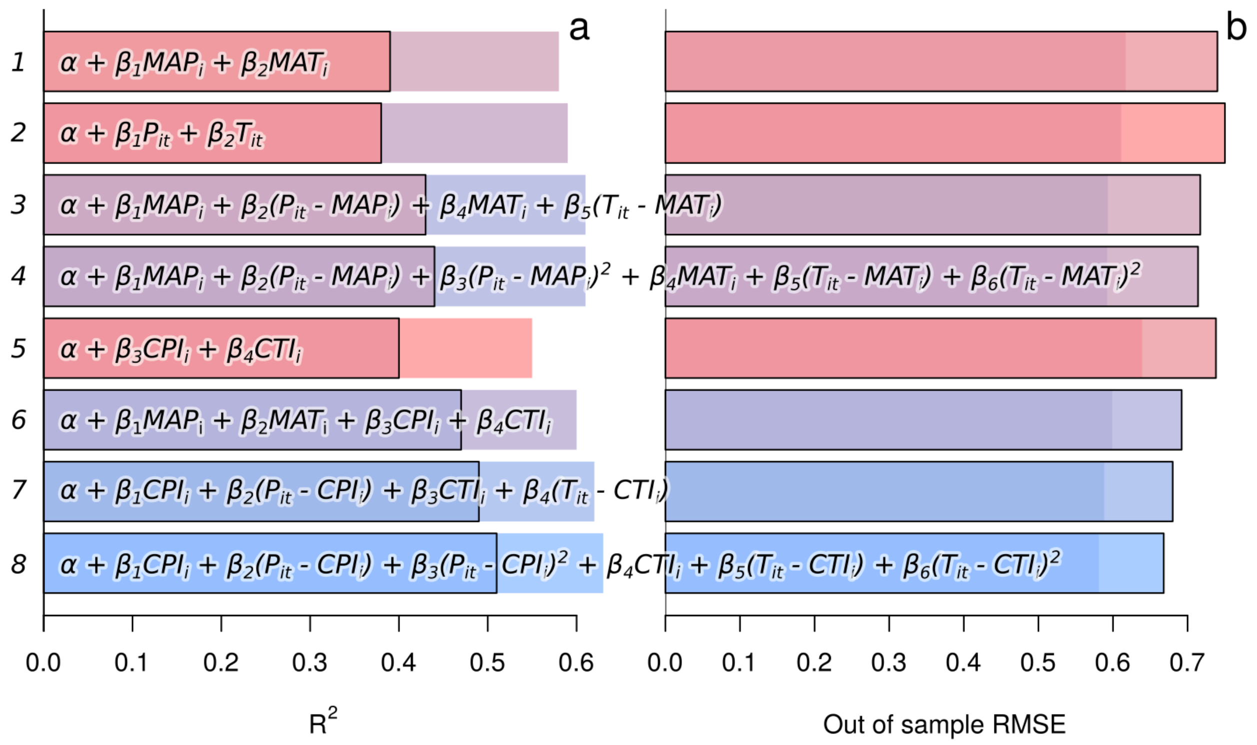


**Figure S1.4.** Results of model comparison for a suite of eight models. The first four models use climatological averages for spatial regressions and as baselines against which to calculate annual anomalies. The latter 4 models allow for disequilibrium by using community climate indices for spatial regressions and to calculate annual disequilibria instead of anomalies. Two versions of each model were evaluated: one with fixed effect predictors only (whose results are shown with outlined bars), and a second with an added smooth term over site coordinates to account for spatial autocorrelation (bars without black outlines). Blue bars indicate the best-supported models, and red bars the least-supported models. Model were compared used two metrics: within-sample coefficient of determination (a), and root-mean-squared-error of the model against out-of-sample test data held out through 5-fold cross-validation (b).

**Supplement 2. Ecoregions analysis**

To gain deeper understanding of the biogeography of disequilibrium and test the robustness of our findings across different ecological contexts, we classified AIM sites according to USEPA level 3 ecoregions. These ecoregions partition the contiguous United States into 85 zones that are determined by shared biotic and abiotic characteristics. The AIM dataset included sites in 31 of these regions. The following ecoregions were most represented in the AIM dataset (Figure S2.1): Northern Basin and Range, Central Basin and Range, Wyoming Basin, Colorado Plateaus, Northwest Great Plains, Middle Rockies, and Mojave Basin and Range. For subsequent analyses, we excluded ecoregions with fewer than 1,000 AIM sites, leaving 13 regions.

Ecoregions varied in their levels of disequilibrium, but most conformed to the broader pattern of negative precipitation and temperature disequilibrium. These ecoregions had the most extreme negative precipitation disequilibria (Figure S2.2, top panel): Northwest Great Plains, Northwest Glaciated Plains, and Snake River Plain. These had the least precipitation disequilibria: Middle Rockies, Colorado Plateaus. These ecoregions had the most extreme negative temperature disequilibria (Figure S2.2, bottom panel): Middle Rockies, Southern Rockies. These had the most positive temperature disequilibria: Sonoran Basin and Range. These had the least temperature disequilibria: Mojave Basin and Range, Northern Basin and Range.

We wished to determine whether our overall findings depended on ecological context beyond that captured by the climate and community composition variables included in our models. To test this, we performed two sets of model fitting where we applied the best-supported model from the overall analysis (Eq. 1) to subsets of the dataset and compared their parameter estimates and predictions. First, we fit the model to data from each ecoregion separately. Second, we fit the model to data from all sites excluding a given ecoregion. We maintained the same spatial random effect structure as in the global model to account for spatial autocorrelation.

As expected, we found substantial variation in ecoregion-specific parameter estimates (Figure S2.2). Parameters pertaining to precipitation effects varied in magnitude, but maintained the same sign across all ecoregions. Temperature effects were more heterogeneous in magnitude and sign, but the most data-rich ecoregions matched the sign of the overall result. Predictions from the models showed a negative effect of extreme disequilibria in nearly all ecoregions, indicating that our overall findings are robust across ecological contexts. Parameter estimates and predictions from datasets with single ecoregions excluded matched the global results closely, indicating that our findings are not dependent on the influence of any one ecoregion (Figure S2.4). Overall, this analysis shows that, had we been limited to a more local dataset, we would have been unlikely to draw a qualitatively different conclusion about the effect of disequilibrium on productivity.

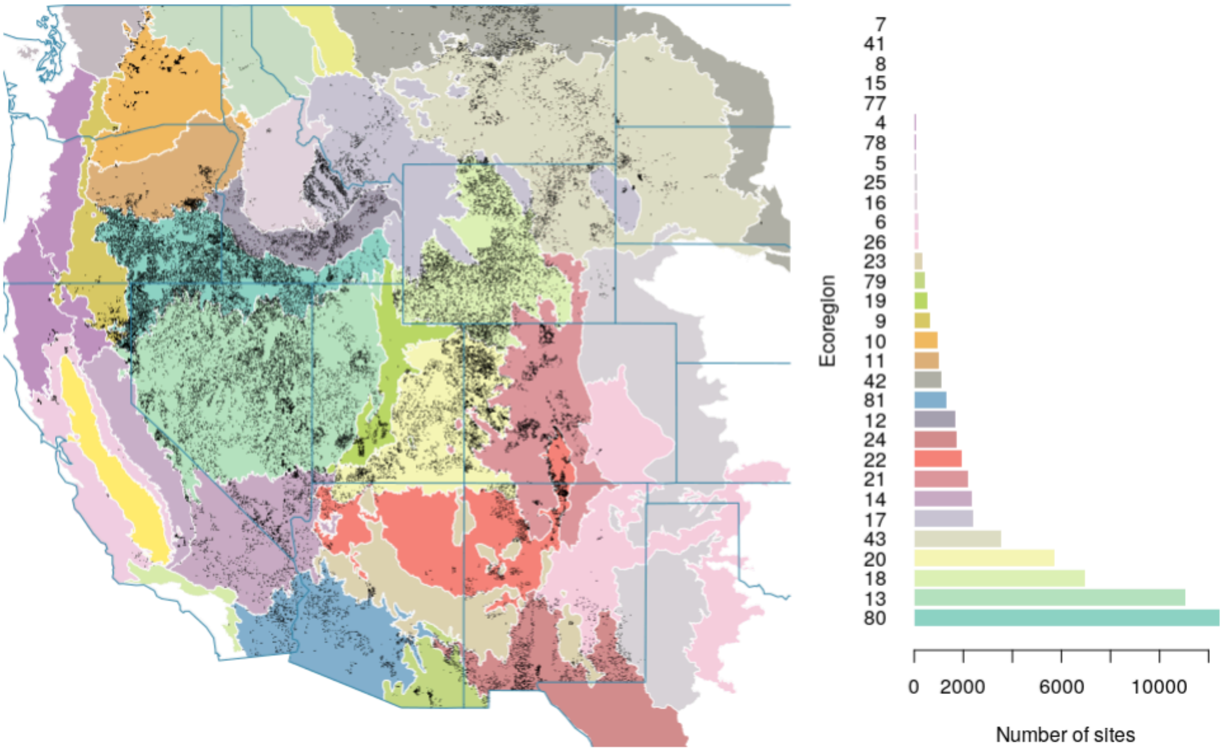


**Figure S2.1.** Ecoregions and represented by the AIM dataset and the number of sites per ecoregion. For the analyses in this supplement, we excluded data from ecoregions with fewer than 1000 sites, removing all beyond ecoregion #42.

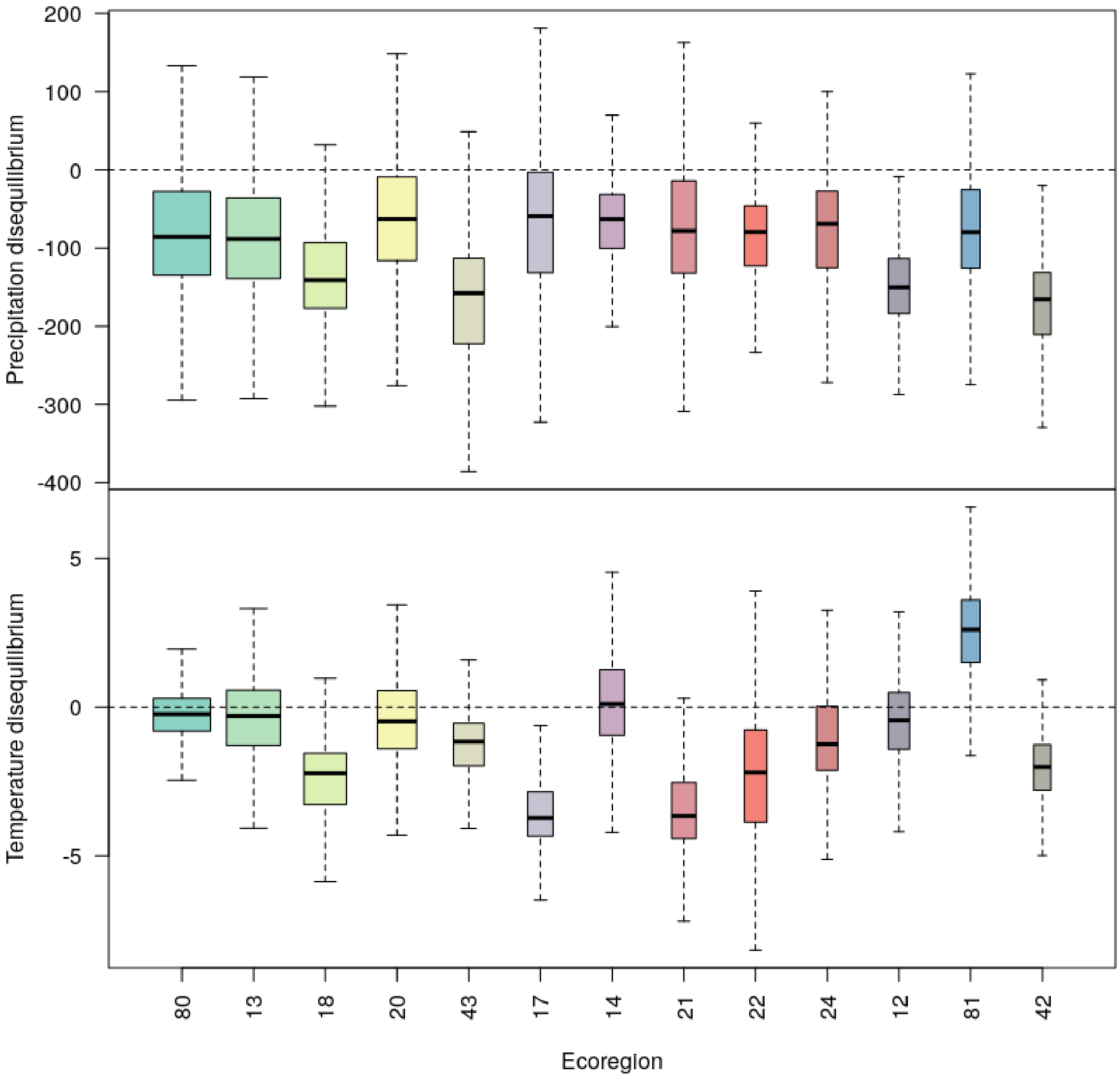


**Figure S2.2.** Precipitation and temperature disequilibria distributions within the ecoregions.

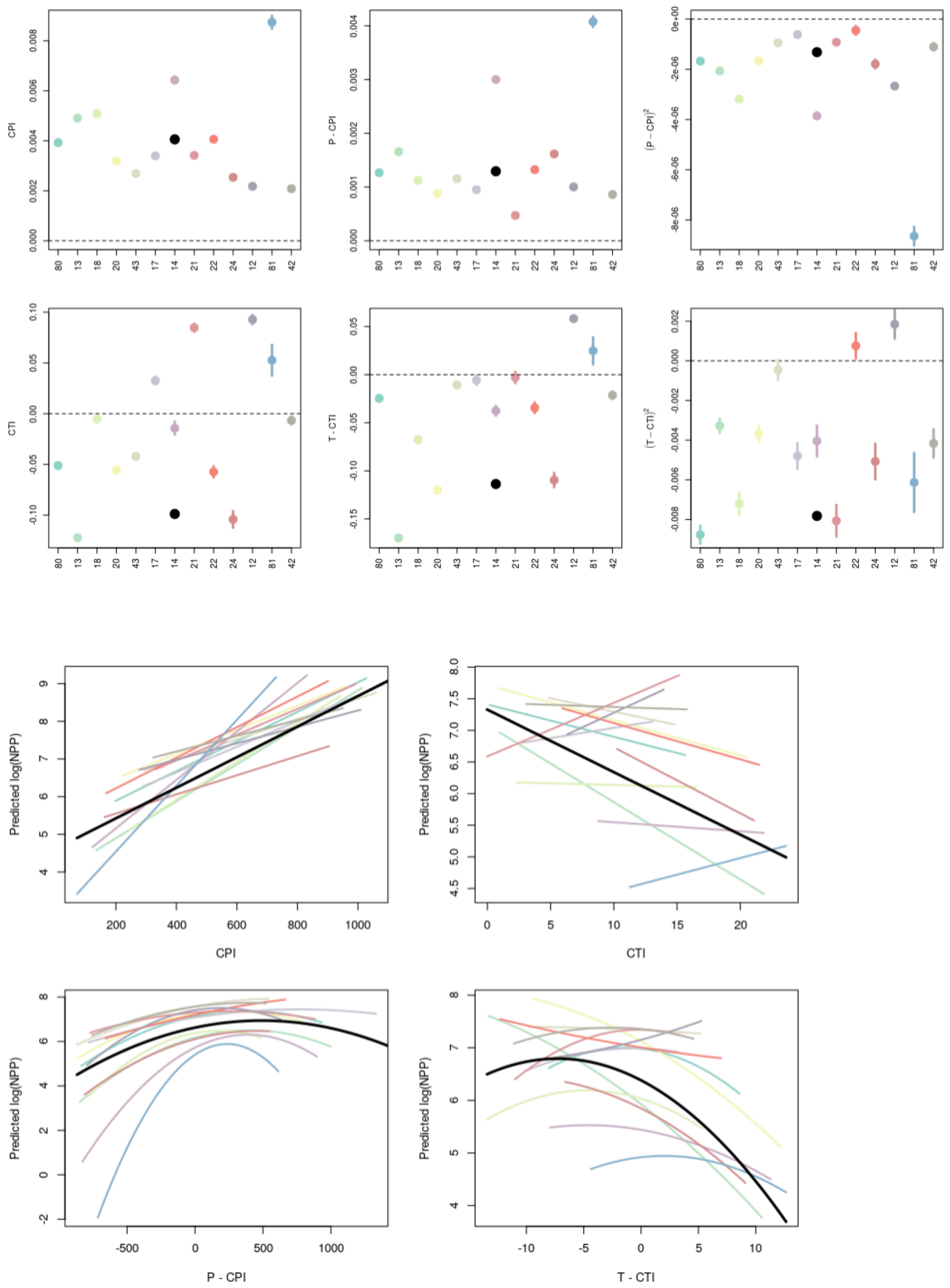


**Figure S2.3.** Parameter estimates and conditional predictions from models fit to data from each ecoregion alone. Colors correspond to figure S2.1.

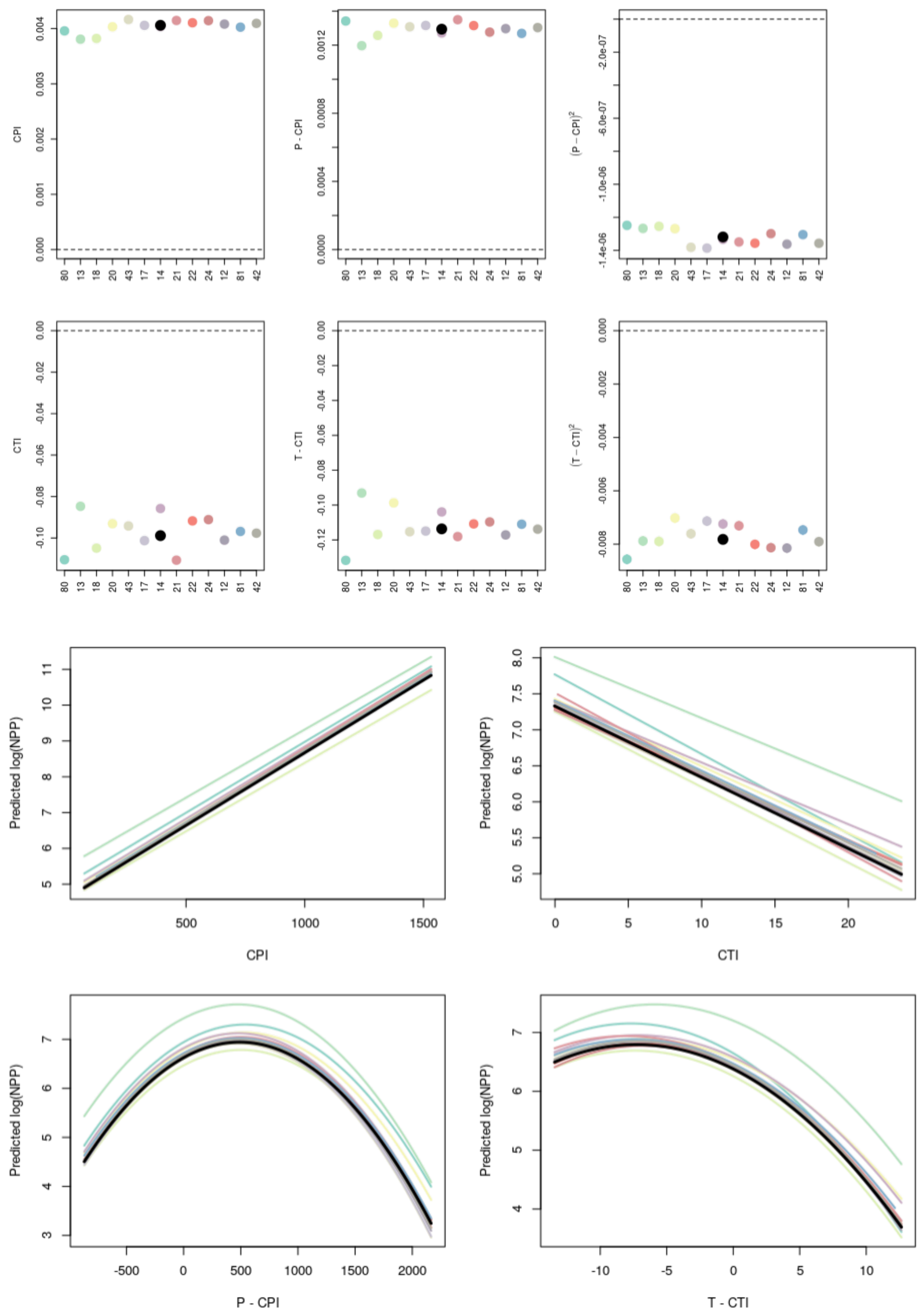


**Figure S2.4.** Parameter estimates and conditional predictions from models fit to data excluding each ecoregion in turn. Colors correspond to figure S2.1, with the color representing which ecoregion was excluded.

**Supplement 3. Alternative modeling approach**

The set of models (Figure 3) used to understand the effect of disequilibrium is appealing because it makes direct comparisons between climatological averages and community climate niches as the baselines against which to judge climate variation. However, the functional form that we used mixes together the direct effects of climate variation and mediating effects of disequilibrium. Specifically, the parameters associated with interannually-varying disequilibrium terms ($(C\text{-}E)^2$) reflect variation in both C and E. No single parameter captures the effect of disequilibrium in isolation, so testing the hypothesis that disequilibrium affects NPP had to be done through model selection rather than by looking at parameter estimates and their uncertainties.

Other modeling approaches might more cleanly separate the equilibrium climate sensitivity of NPP from the effect of disequilibrium that modifies that relationship. We turned to our recent theoretical work (Stemkovski et al. 2026) to explore other functional forms. One option from that study modifies the equilibrium sensitivity of ecosystem function with a gaussian disequilibrium term.

$$F_t = (\alpha + \beta C_t)e^{-(C_t - E_t)^2/\gamma}$$

In this equation, ecosystem function *F* is predicted by climate *C* and the community climate niche *E*. The parameters alpha and beta represent the equilibrium sensitivity, while the parameter gamma controls the effect of disequilibrium. This functional form is inspired by a global plant production model that incorporates the effect of lagging physiological acclimation to temperature for photosynthetic efficiency into estimates of global NPP (Friend 2010). disequilibrium (Figure S3.1).

We applied this modeling approach to the present dataset with this form that separates the equilibrium relationship *f* between NPP and climate from the disequilibrium effect *g*:

$$log(NPP_{it}) = f(P_{it}, T_{it}, MAP_i, MAT_i) * g(P_{it}, T_{it}, CPI_i, CTI_i)$$

Here, *f* is the pattern that would be expected if disequilibrium was always zero. It is a function of interannually varying precipitation and temperature and long-term site climatology.

$$f = \alpha + \beta_{1P}MAP_i + \beta_{2P}(P_{it} - MAP_i) + \beta_{1T}MAT_i + \beta_{2T}(T_{it} - MAT_i)$$

Other models of the equilibrium sensitivity (perhaps with a saturating precipitation relationship) might be more appropriate. The effect of disequilibrium *g* modifies NPP when disequilibrium is large:

$$g = exp\left(\frac{-(P_{it}-CPI_i)^2}{\gamma_P}\right) * exp\left(\frac{-(T_{it}-CTI_i)^2}{\gamma_T}\right)$$

Here, we used two separate disequilibrium terms for temperature and precipitation disequilibrium. The effects of these disequilibria are controlled by the gamma parameters.

We fit this model to the AIM and RAP dataset using the *nlsLM* function from the *minpack.lm* package. We did not account for spatial autocorrelation in this analysis, but this should be done to ensure robust inference. We found that both temperature and precipitation disequilibria significantly reduced NPP relative to what would be expected if communities were in equilibrium with climate (Figure S3.2). The inference from this model differed somewhat from the model in the main analysis. The results of the main model suggested that NPP is maximized when communities receive more precipitation than what they are suited to, and the alternative model showed an even stronger positive effect of increased precipitation, with peak NPP further from community-climate equilibrium (Figure S3.3 left panel). The alternative model also predicted a stronger negative relationship between temperature and productivity, but it agreed with the main model that both negative and positive temperature disequilibria reduce productivity (Figure 3.3 right panel). The alternative model formulation necessarily bounds predicted log(NPP) above zero, which prevents unrealistic extrapolations outside of the data range which can be seen in Figure 5.

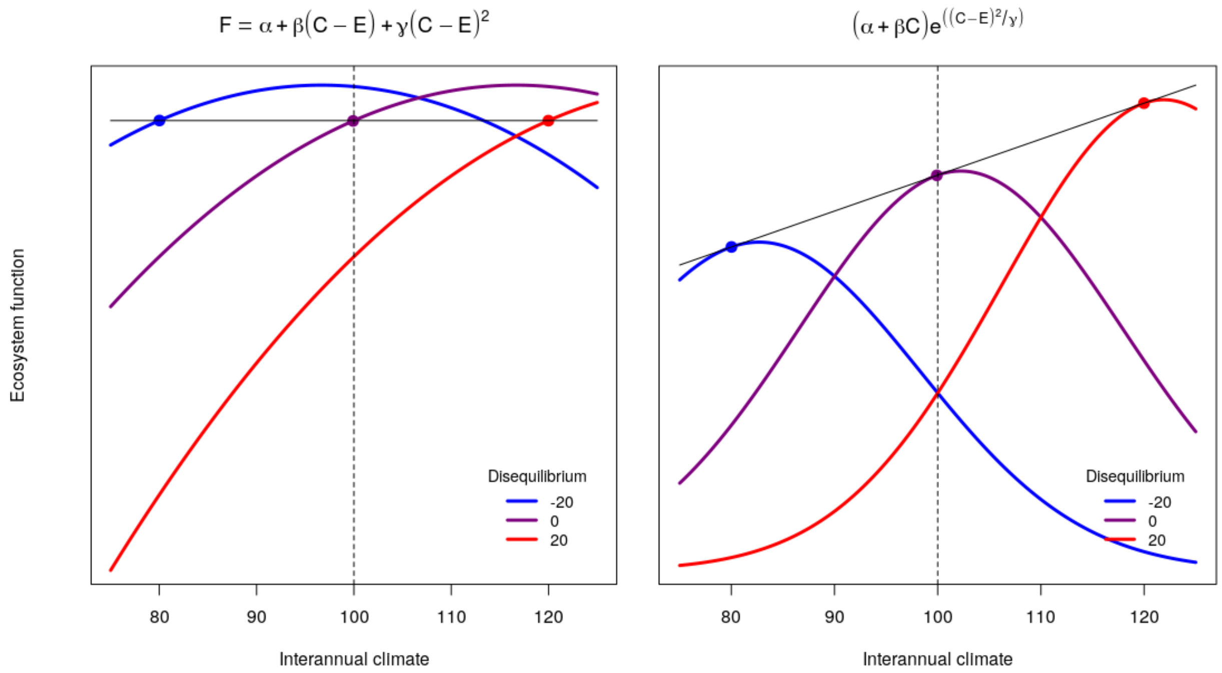


**Figure 3.1.** The present modeling framework conflates the direct effect of interannual climate variation and the effect of disequilibrium (left panel). An alternative model would explicitly separate equilibrium sensitivity from the effect of disequilibrium (right panel). The colored curves show predicted ecosystem function for three hypothetical communities experiencing the same climate but having different disequilibrium values. The black solid lines show the equilibrium sensitivity when the disequilibrium terms are zero (*E*=*C*). Modified from Stemkovski et al. (*in press*).

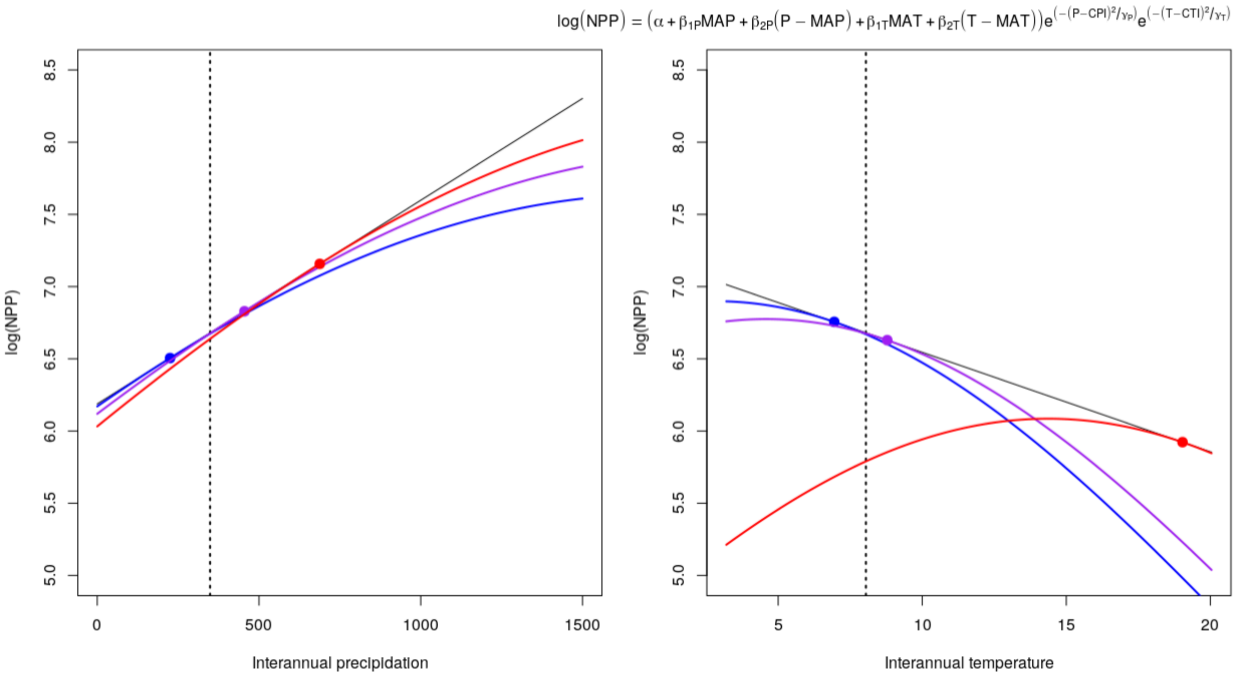


**Figure 3.2.** Disequilibrium causes ecosystem function to deviate from the equilibrium expectation. The colored curves show predicted NPP for cross-sections of community climate niche values, with the 2.5% (blue), 50% (purple) and 97.5% (red) quantiles of the total distributions across sites. The black solid lines show the equilibrium sensitivity when the disequilibrium terms are always zero (*CPI*=*P* & *CTI*=*T*). The vertical dashed lines show the means of the climate variables across sites.

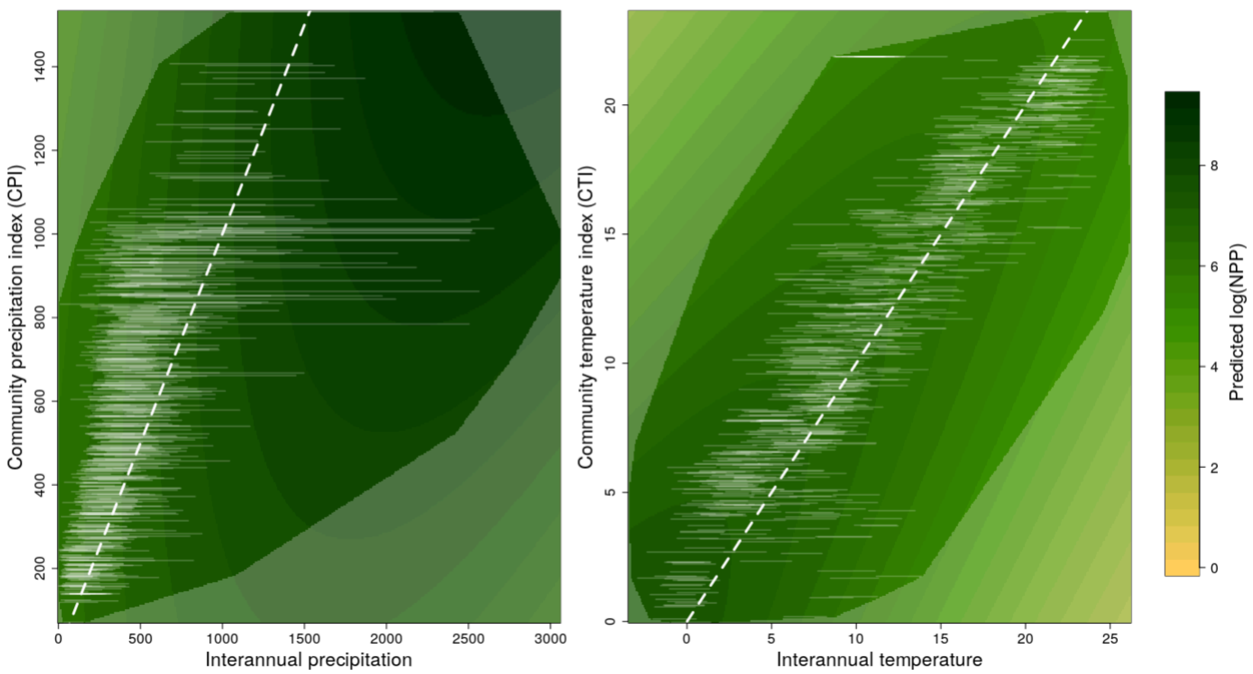


**Figure 3.3.** Predictions over climate and community climate niche space from the nonlinear model. The symbols in this figure are the same as in Figure 5.

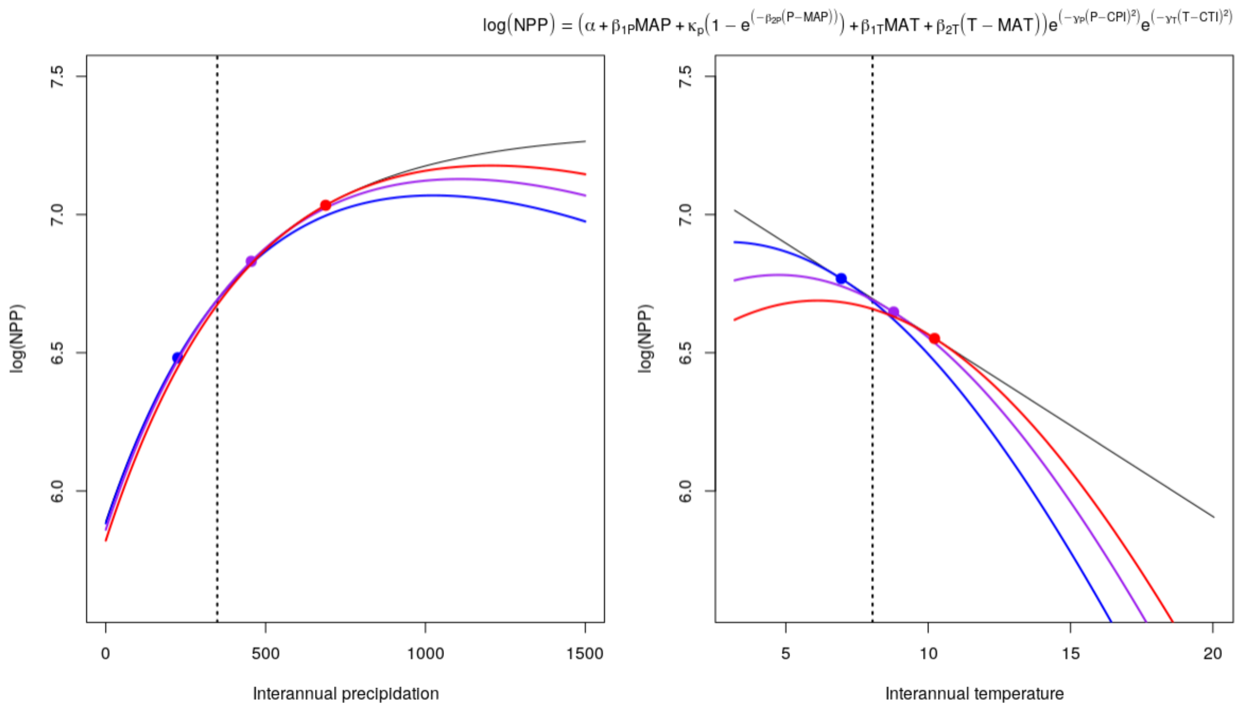


**Figure 3.4.** Predictions from a version of the model with nonlinear equilibrium sensitivity to interannual precipitation variation.

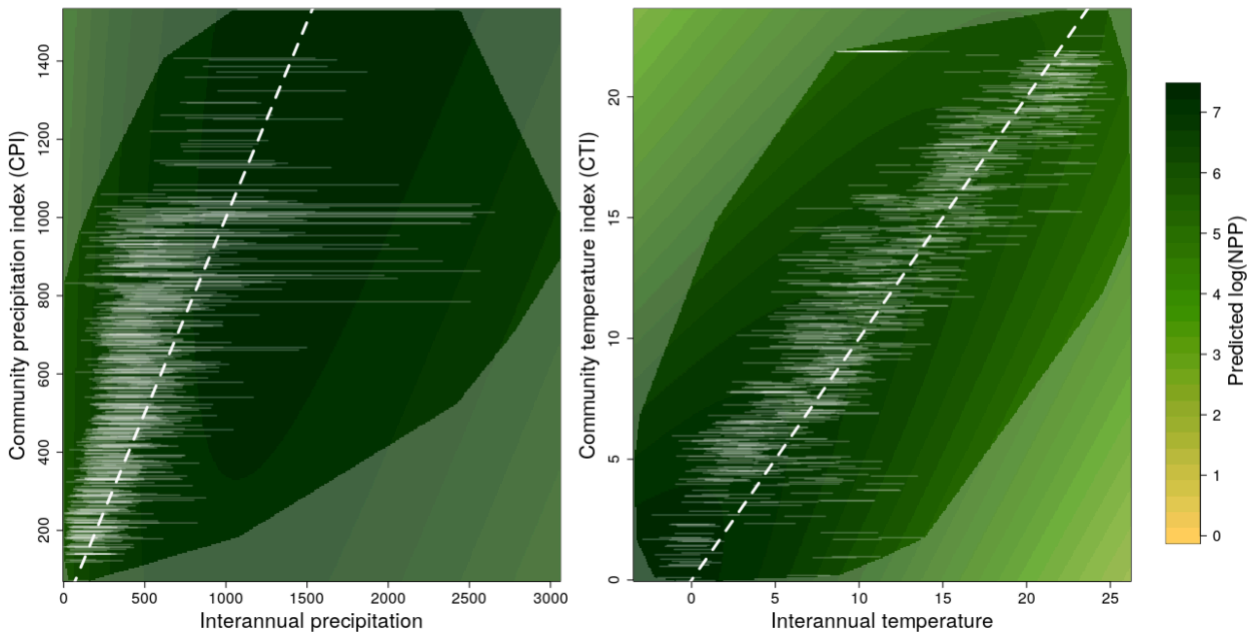


**Figure 3.5.** Predictions from a version of the model with nonlinear equilibrium sensitivity to interannual precipitation variation.

### Supplement 4. Additional analyses

#### 4.1 Temporal shifts in community climate indices

Because sites in the AIM dataset were not resurveyed, we were not able to directly assess whether community climate niches changed over time or whether they varied interannually due to weather variation within sites. However, we took advantage of variation in survey years (2010-2024) to examine temporal trends over time across sites. We found that community temperature niches became 0.027°C warmer and 1.72mm wetter per year, though time explained <1% of the variation in CTI and CPI across sites. These trends indicate ongoing thermophilization and hydrophilization of rangeland communities – shifts that could exacerbate current disequilibrium. Both rates were statistically significant, but were much slower than would be needed to substantially affect the magnitude of community-climate disequilibrium over the study period.
